# Photo-induced changes in tissue stiffness alter epithelial budding morphogenesis in the embryonic lung

**DOI:** 10.1101/2024.08.22.609268

**Authors:** Kara E. Peak, Poornacharanya Rajaguru, Adil Khan, Jason P. Gleghorn, Girgis Obaid, Jacopo Ferruzzi, Victor D. Varner

## Abstract

Extracellular matrix (ECM) stiffness has been shown to influence the differentiation of progenitor cells in culture, but a lack of tools to perturb the mechanical properties within intact embryonic organs has made it difficult to determine how changes in tissue stiffness influence organ patterning and morphogenesis. Photocrosslinking of the ECM has been successfully used to stiffen soft tissues, such as the cornea and skin, which are optically accessible, but this technique has not yet been applied to developing embryos. Here, we use photocrosslinking with Rose Bengal (RB) to locally and ectopically stiffen the pulmonary mesenchyme of explanted embryonic lungs cultured *ex vivo*. This change in mechanical properties was sufficient to suppress FGF-10-mediated budding morphogenesis along the embryonic airway, without negatively impacting patterns of cell proliferation or apoptosis. A computational model of airway branching was used to determine that FGF-10-induced buds form via a growth-induced buckling mechanism and that increased mesenchymal stiffness is sufficient to inhibit epithelial buckling. Taken together, our data demonstrate that photocrosslinking can be used to create regional differences in mechanical properties within intact embryonic organs and that these differences influence epithelial morphogenesis and patterning. Further, this photocrosslinking assay can be readily adapted to other developing tissues and model systems.

## Introduction

Mechanical inputs, such as forces, deformations, and changes in stiffness, have been shown to influence aspects of embryonic development^1,2^. Time-varying changes in mechanical properties, for instance, are associated with a variety of developmental events, including early embryonic cleavage, notochord elongation, gastrulation, and cardiac looping^3–7^. This work has revealed that differences in mechanics are present at different embryonic stages, but there is little *in vivo* (or *ex vivo*) data demonstrating that altered mechanical properties can directly influence organ patterning and morphogenesis. A variety of *in vitro* assays have been used to show that changes in substratum stiffness can regulate the differentiation of isolated progenitor cells^8^, but this idea has yet to be demonstrated within the context of organogenesis, in part owing to a lack of experimental tools to modulate mechanical properties within intact embryonic organs. Recent work has shown that the extracellular matrix (ECM) degradation or patterning can influence branching morphogenesis^9,10^, but these changes have not been connected directly to changes in mechanical properties. Here, we present a new technique that uses photocrosslinking to locally modulate tissue stiffness in embryonic lungs cultured *ex vivo* and assess how these mechanical changes influence epithelial budding.

Exposure to light can elicit a variety of specific chemical reactions, which over the years have proved useful for several biomedical applications^11^. Photosensitive synthetic polymers, for instance, have been used to create hydrogel-based biomaterials with tunable mechanical properties for 3D cell and organoid culture^12–14^. In addition, the light-initiated crosslinking of native ECM with specific photo-activatable dyes, such as riboflavin and Rose Bengal (RB), has been used to stiffen collagen gels^15,16^, restore mechanical integrity to diseased corneas^17–20^, and aid in dermal wound closure^21^. To our knowledge, however, these photocrosslinking assays have never been adapted to the study of embryonic organs, possibly owing to the need for tissues to be optically accessible. RB is activated by the presence of green light (530-560 nm)^22^ and can alter ECM crosslinking and mechanical properties within both 3D collagen gels^15^ and soft tissues like the cornea and skin^19,21^ (Fig. 1A). Here, we capitalized on the biocompatibility of RB and adapted RB-mediated photocrosslinking to the *ex vivo* culture of an explanted embryonic organ, using the developing avian lung as an illustrative example.

**Figure 1:**
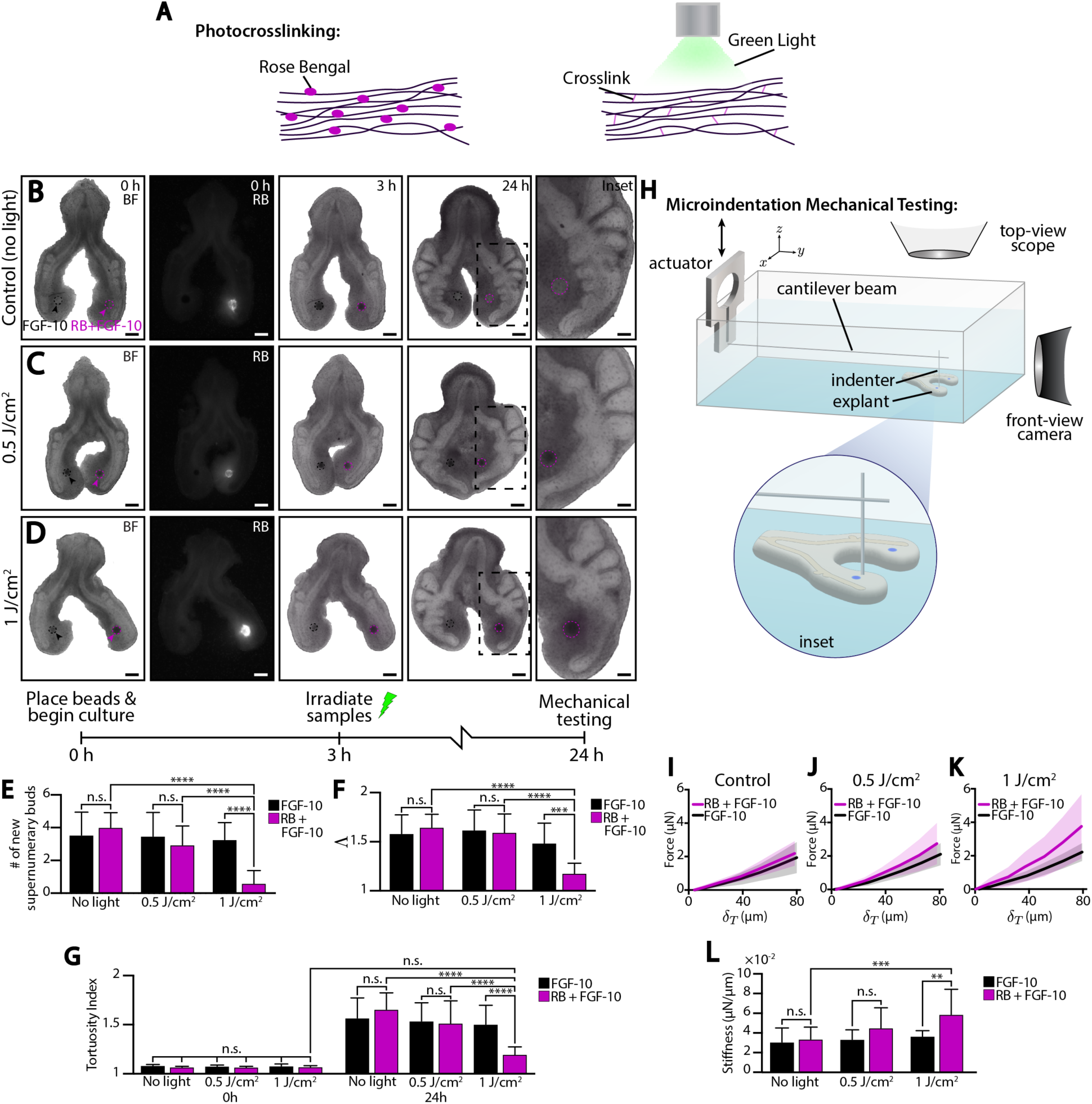
Photocrosslinking leads to increased tissue stiffness and inhibits FGF-10-induced epithelial buckling. (A) Schematic representation of Rose Bengal (RB), shown as magenta ovals, associating with extracellular matrix (ECM) proteins, shown as black strands. Exposure to green light leads to activation of Rose Bengal, resulting in the formation of crosslinks, shown as magenta lines, between ECM proteins. (B-D) Bright-field and fluorescent images of representative embryonic lung explants cultured ex vivo with either FGF-10 beads (left lobe, black arrow and dashed lines) or FGF-10 + RB beads (right lobe, magenta arrow and dashed lines). Fluorescent images show RB autofluorescence visualized by green light. Cultured lungs were mechanically tested at the end of 24 h. Scale bars, 200 µm. (Inset scale bars, 100 µm). (E-G) Quantification of (E) number of new supernumerary buds, (F) the epithelial contour ratio Λ = *L*⁄*L*_0_, and (G) the tortuosity index 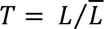 where *L* is the measured length of the ventral epithelium at each time-point, 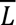 is the length of the straight line connecting the endpoints of *L*, and *L*_5_ is the length of the ventral epithelium at t = 0 h, which is used as a reference length. A one-way ANOVA (E-F) or two-way ANOVA (G), followed by a Tukey post-hoc test, were used to make statistical comparisons. (Control, FGF-10: n = 15; Control, RB + FGF-10: n = 13; 0.5 J/cm^2^, FGF-10: n = 15; 0.5 J/cm^2^, RB + FGF-10: n = 15; 1 J/cm^2^, FGF-10: n = 12; 1 J/cm^2^, RB + FGF-10: n = 12; * p < 0.05, ** p < 0.01, *** p < 0.001, **** p < 0.0001; error bars represent s.d.). (H) Schematic representation of microindentation mechanical testing. (I-K) Averaged force versus tissue deflection (*δ_T_*) curves for (I) control, (J) 0.5 J/cm^2^, and (K) 1 J/cm^2^. Curves represent the mean value while the shaded region represents standard deviation. (L) Stiffness plot for all treatment groups. A two-way ANOVA, followed by a Tukey post-hoc test, was used to make statistical comparisons. (Control, FGF-10: n = 15; Control, RB + FGF-10: n = 15; 0.5 J/cm^2^, FGF-10: n = 14; 0.5 J/cm^2^, RB + FGF-10: n = 13; 1 J/cm^2^, FGF-10: n = 12; 1 J/cm^2^, RB + FGF-10: n = 11; * p < 0.05, ** p < 0.01, *** p < 0.001, **** p < 0.0001; error bars represent s.d.).

Numerous organs, including the lung, kidney, and salivary gland, form via a process known as branching morphogenesis, wherein a simple epithelial tube undergoes a series of branching events to build a ramified tree^23,24^. In the embryonic lung, airway branching involves distinct branching modes: lateral branches, which emerge along the length of a parent branch, and bifurcations, in which the tip of a parent branch splits^25^. This process is regulated, in part, by reciprocal signaling interactions between the branching airway epithelium and the surrounding layer of pulmonary mesenchyme^26^, but branching morphogenesis can also be influenced by changes in mechanics^27–35^. Focal regions of fibroblast growth factor (FGF)-10 expression in the pulmonary mesenchyme are thought to specify the formation of individual buds and provide a biochemical template for the overall branching pattern^36^. However, we have recently shown that individual sources of FGF-10 can promote bud formation via epithelial buckling^37^. In general, buckling morphogenesis can depend on relative differences in mechanical properties between adjacent tissue layers^28,38^; however, previous studies have not yet explored how altered tissue stiffness influences FGF-10-induced budding. Here, we present a new RB photocrosslinking assay to locally and ectopically stiffen the pulmonary mesenchyme of cultured embryonic avian lungs. This technique is then combined with micro-mechanical testing, quantitative fluorescence microscopy, and computational modeling to determine how regional differences in stiffness influence FGF-10-induced budding morphogenesis.

## Results

### Ectopically stiffening the pulmonary mesenchyme inhibits FGF-10-induced budding morphogenesis

We embedded RB-loaded beads within the pulmonary mesenchyme of explanted embryonic avian lungs, irradiated them with green light (fluence levels of 0, 0.5, or 1 J/cm^2^) to promote photocrosslinking of the tissue, and determined how changes in mechanical properties influence FGF-10-mediated budding morphogenesis. In previous work, we showed that individual focal sources of FGF-10 can elicit the formation of multiple supernumerary buds along the primary bronchus of cultured explants^37^. Following irradiation with light, agarose beads loaded with both RB and FGF-10 showed decreased levels of bud formation at increased fluence levels, as compared to either “no light” controls (containing RB) or contralateral controls (containing only FGF-10) (Fig. 1B-E). These data suggested that increased levels of RB-mediated photocrosslinking were suppressing the formation of FGF-10-induced supernumerary buds. (Higher fluence levels (≥ 2 J/cm^2^) compromised the viability of the cultured explants (Supplementary Fig.1)). Morphometric analysis of epithelial geometry further highlighted these differences. Treatment with FGF-10 alone elicited an increase in both epithelial length and tortuosity, which coincided with the emergence of new epithelial buds (Fig. 1E-G); however, RB-mediated photocrosslinking (at 1 J/cm^2^) produced a less elongated and less tortuous epithelium with fewer FGF-10-induced buds as compared to both “no light” and contralateral controls (Fig. 1F-G).

To confirm that photocrosslinking with RB produced a change in tissue stiffness, we used our customized microindentation system to quantify regional differences in the mechanical properties of the pulmonary mesenchyme (Fig. 1H; Supplementary Fig. 2). Irradiated lungs exhibited a fluence-dependent increase in mesenchymal stiffness, wherein the lobes containing an RB-loaded bead were significantly stiffer than either “no light” or contralateral controls (Fig. 1I-K). Stiffness was evaluated by computing the local force-deflection curve, with an increased slope indicating a stiffer tissue. At the highest fluence level (1 J/cm^2^), we observed the largest increase in tissue stiffness (Fig. 1L), while no difference in mechanical properties was observed between RB- and non-RB-treated lobes in the absence of light (Fig. 1L). Importantly, these microindentation results were consistent with the increase in stiffness observed among photocrosslinked 3D collagen gels upon treatment with RB (Supplementary Fig. 3). Taken together, these data suggest that RB-mediated photocrosslinking can locally alter the mechanical properties of the developing pulmonary mesenchyme and suppress the formation of FGF-10-induced buds.

### Treatment with RB does not adversely affect the patterns of proliferation and apoptosis associated with epithelial budding

Previous work has shown that the formation of FGF-10-induced supernumerary buds is associated within an increase in epithelial proliferation^37^. To confirm that decreased budding in RB-treated lungs was not accompanied by lower levels of proliferation, we quantified patterns of EdU incorporation along the airway epithelium of both photocrosslinked and non-photocrosslinked lung explants (Fig. 2A-C). We did not observe a significant reduction in the number of EdU-positive nuclei following irradiation with green light nor upon treatment with RB (Fig. 2D). In fact, at the highest fluence level (1 J/cm^2^), in which we observed a decrease in FGF-10-induced budding (Fig. 1), we observed a modest increase in epithelial proliferation (Fig. 2D); however, the increased stiffness associated with this treatment condition was sufficient to inhibit the formation of any supernumerary buds (Fig. 1E, 2C-D).

**Figure 2:**
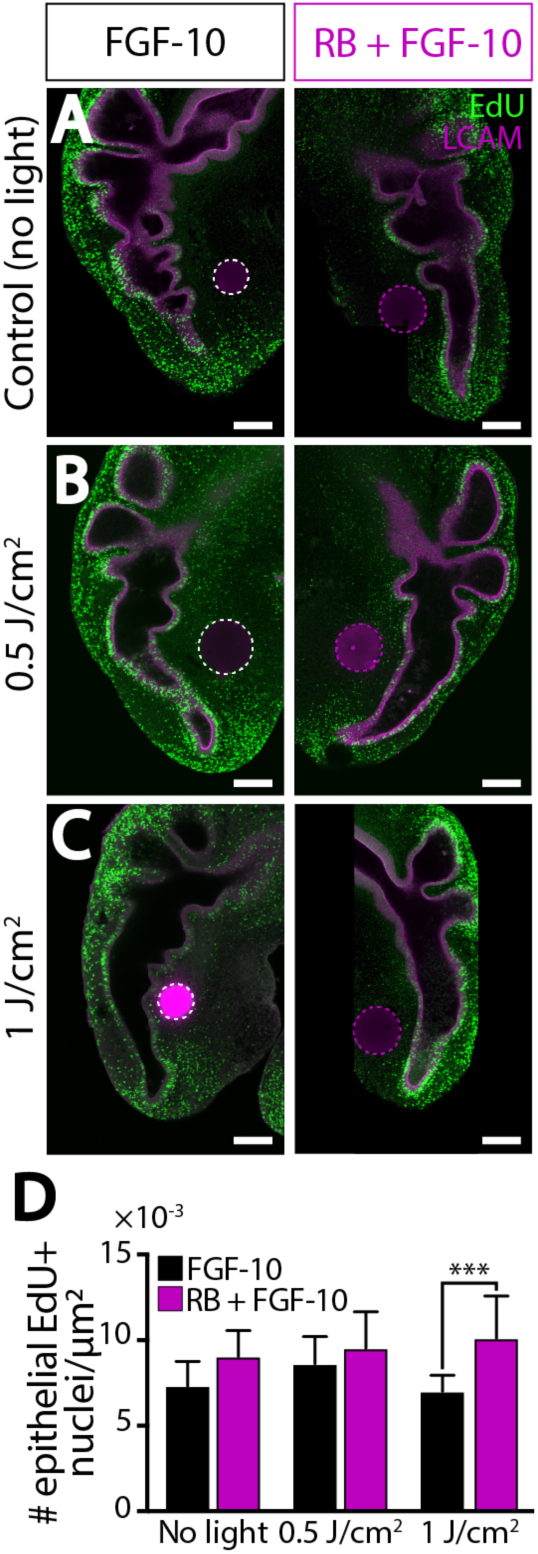
Photocrosslinking does not reduce rates of proliferation along the embryonic airway epithelium. (A-C) Confocal images of EdU incorporation and LCAM immunofluorescence in representative whole-mount lung explants. White or magenta dashed lines denote FGF-10 or RB + FGF-10-soaked beads, respectively. Scale bars, 200 μm. (D) Quantification of EdU incorporation within the ventral epithelium. A one-way ANOVA, followed by a Tukey post-hoc test, was used to determine significance between groups (Control, FGF-10: n = 12; Control, RB + FGF-10: n = 13; 0.5 J/cm^2^, FGF-10: n = 13; 0.5 J/cm^2^, RB + FGF-10: n = 12; 1 J/cm^2^, FGF-10: n = 13; 1 J/cm^2^, RB + FGF-10: n = 12; * p < 0.05, ** p < 0.01, *** p < 0.001, **** p < 0.0001; error bars represent s.d.).

Previous work has suggested that RB-mediated crosslinking in the cornea can be associated with increased cell death at high fluence levels^17^. To determine if decreased epithelial budding in RB-treated lungs was associated with a change in apoptosis, we stained cultured explants for CC3 immunofluorescence. Neither irradiation with green light nor treatment with RB appeared to cause any qualitative changes in cell apoptosis. We did not observe elevated levels of CC3-positive nuclei within the airway epithelium of ectopically stiffened lungs, as compared to either “no light” or contralateral controls (Fig. 3A-B, Supplementary Fig. 4A-D). However, as a positive control, explants cultured in the presence of staurosporine, a pharmacological activator of apoptosis, elicited increased numbers of CC3-positive nuclei within all treatment conditions (Fig. 3C-D, Supplementary Fig. 4E-G). Thus, neither decreased proliferation nor increased apoptosis appear to be associated with the changes in FGF-10-induced bud formation that follow RB-mediated photocrosslinking.

**Figure 3:**
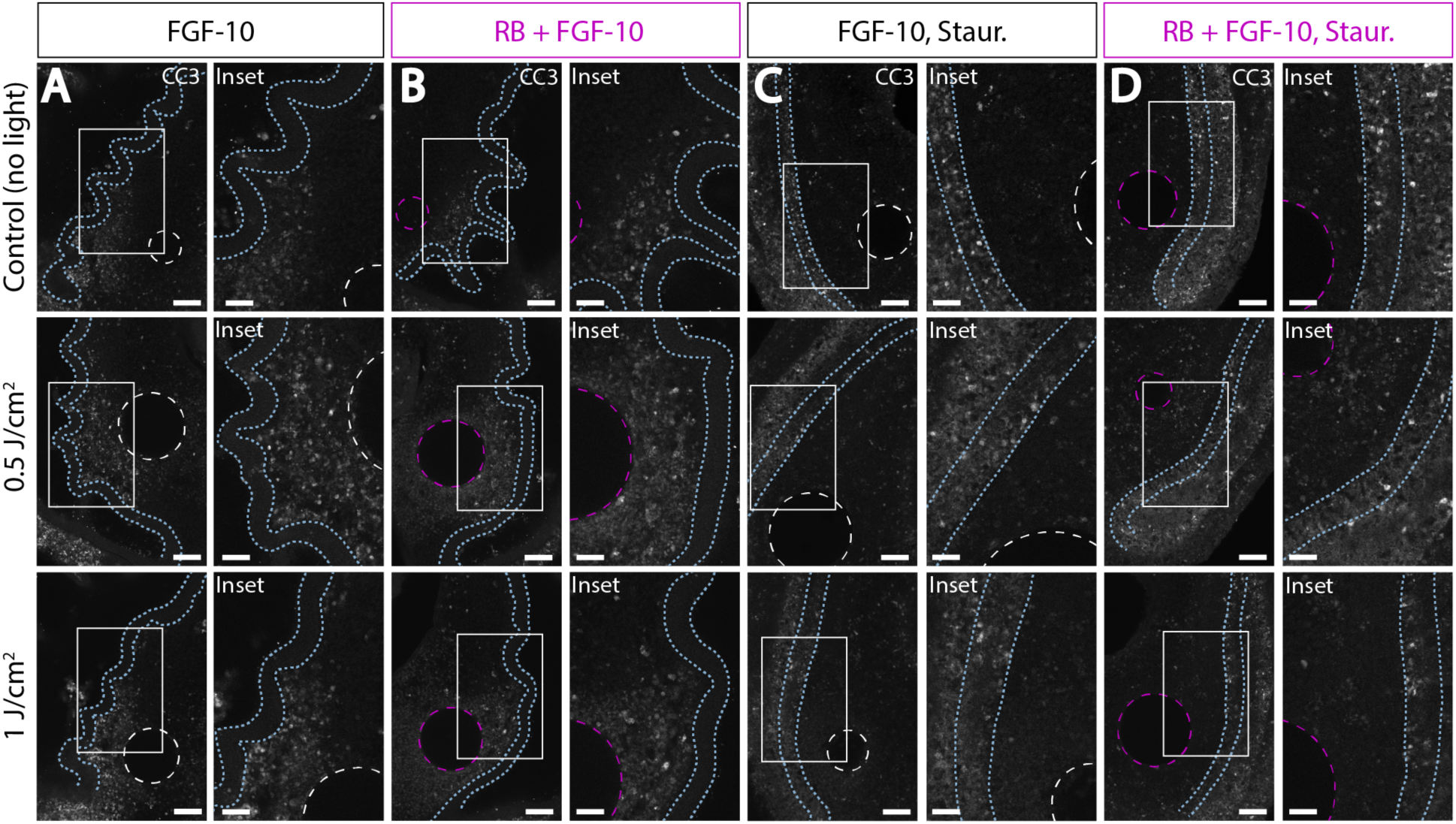
No apparent increase in apoptosis in photocrosslinked lung explants. (A-D) Confocal images of CC3 immunofluorescence in representative whole-mount lung explants. Some explants were treated with staurosporine (staur.), an activator of cell apoptosis (C-D). White or magenta dashed circles denote FGF-10 or RB + FGF-10-soaked beads, respectively. Blue dashed lines outline the ventral epithelium. Inset region indicated with white box. Scale bars, 100 μm. Insets scale bars, 50 μm. (Control: n = 9; 0.5 J/cm^2^: n = 9; 1 J/cm^2^: n = 9; Control, staurosporine: n = 9; 0.5 J/cm^2^, staurosporine: n = 9; 1 J/cm^2^, staurosporine: n = 9).

### Computational model of FGF-10-induced budding morphogenesis suggests that increased mesenchymal stiffness can suppress epithelial buckling

To determine how tissue stiffness influences FGF-10-driven budding morphogenesis, we created a computational model of the embryonic airway and surrounding pulmonary mesenchyme that combined morphogen diffusion and continuum mechanics (Fig. 4). As a first approximation, the airway epithelium was modeled as a thin layer of tissue overlying a thick mesenchyme containing a single focal source of FGF-10 (Fig. 4A, Supplementary Fig. 5). As described previously^39^, tissue growth was modeled by decomposing the overall deformation of the tissue (***F***) into a component due to elastic deformation (***F***^∗^) and a component due to growth (***G***), where ***F*** = ***F***^∗^ · ***G*** and growth is specified via the components of the growth tensor ***G***. FGF-10 was allowed to diffuse from this focal source, and the amount of growth within the epithelium (i.e., the components of ***G***) were dependent upon the local concentration of FGF-10, as governed by the growth law 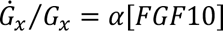, where *α* represents a growth constant, *G*_*x*_ is the longitudinal component of ***G***, [*FGF*10] is the concentration of FGF-10, and 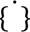 denotes differentiation with respect to time (Fig. 4A, Supplementary Fig. 5). (Additional model details are provided in the Methods.) In addition, since previous studies have suggested a difference in mechanical properties between mesenchymal and epithelial tissues^40,41^, as a first approximation, we assumed the shear modulus of the mesenchyme (*C_m_*) was smaller than that of the epithelium (*C_e_*), and the normalized mesenchymal shear modulus 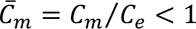. At the beginning of the simulation, no growth was present within either the epithelium or mesenchyme (*G*_*x*_ = 1) (Fig. 4B). However, as time progressed, FGF-10 diffusion promoted longitudinal growth within the epithelium that caused this layer to buckle and create multiple bud-like structures, which formed simultaneously (Fig. 4B, Supplementary Movies 1 and 2) in a manner similar to that observed in cultured embryonic lungs^37^. Before buckling, compressive longitudinal stresses developed within the epithelium as *G*_*x*_ increased, since epithelial expansion was being constrained by the adjacent layer of mesenchyme (Fig. 4B). Once the FGF-10-induced buds had formed, however, the distribution of stresses through the thickness of the epithelium was consistent with bending (i.e., compressive and tensile stresses located along concave and convex regions of the epithelium, respectively)^42^.

**Figure 4:**
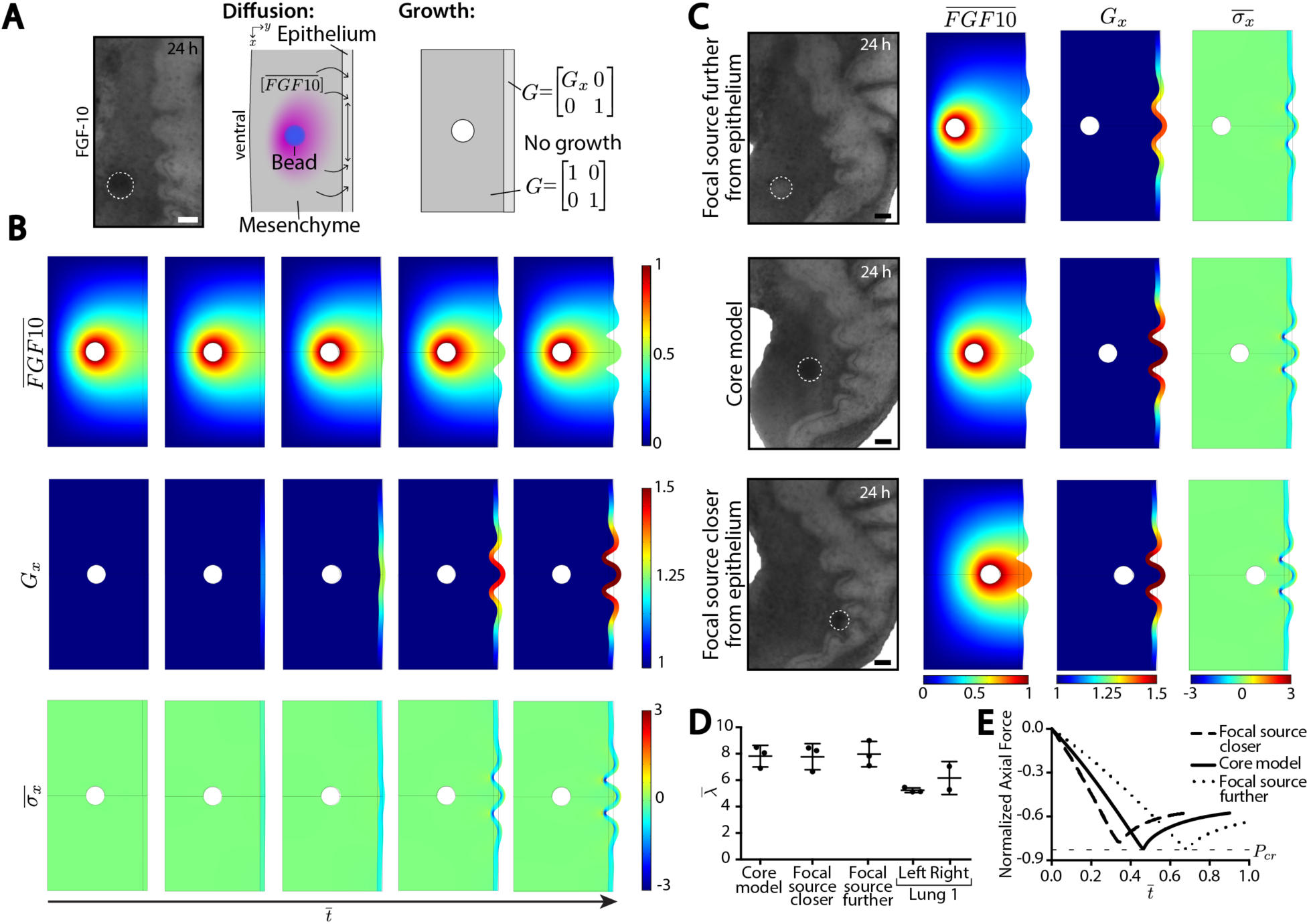
Computational model of FGF-10-driven buckling morphogenesis. (A) Representative bright-field image inset of an embryonic lung explant cultured with an FGF-10-loaded bead and a buckled ventral epithelium. Scale bar, 50 µm. Schematic representation of the cultured lung showing FGF-10 diffusion from the bead and stimulating the ventral epithelium to grow. Schematic representation of the computational model based on the experimental system. Growth (***G***) within the modeled epithelial bar is specified uniaxially (*G_x_*), but the modeled mesenchyme does not grow. (B) Core model results over non-dimensionalized time 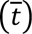 for normalized FGF-10 concentration 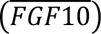, growth in the x-direction (*G_x_*), and normalized Cauchy stress 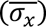. (C) Varying the location of the focal source of growth factor to be further or closer to the epithelium. Representative bright-field images show an FGF-10-loaded bead at various distances from the epithelium. White dashed line indicates bead location. Scale bar = 50 µm. Results for normalized FGF-10 concentration 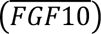, growth in the x-direction (*G_x_*), and normalized Cauchy stress 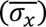 shown at the final timepoint of the models. (D) The normalized wavelength for all three models compared to a representative lung explants shows a characteristic value. (E) Normalized axial force versus non-dimensionalized time 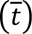 for all three models. All reach a deflection point, interpreted as the critical load (*P_cr_*) and denoted by the horizontal dashed line.

In previous experiments, we observed robust FGF-10-induced bud formation even if the distance between the airway epithelium and the focal source of FGF-10 was varied^37^. To determine how bead placement influences epithelial buckling, we varied the location of FGF-10-loaded bead in our model and assessed its influence on budding morphogenesis (Fig. 4C). Although changes in bead location produced subtle differences in the amplitude of the epithelial folds, we still observed robust buckling in all cases, with FGF-10-induced buds exhibiting the same (normalized) characteristic wavelength 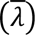, as seen experimentally^37^ (Fig. 4C-D). In addition, each simulation predicted the onset of buckling at the same critical axial load (*P_cr_*), which plateaued following the formation of new epithelial buds (Fig. 4E)^42^. Systematic analysis of other model parameters, such as the magnitude of the negligible transverse point force (Supplementary Fig. 6), the mesh size (Supplementary Fig. 7), the growth constant (*α*) (Supplementary Fig. 8), the nondimensional FGF-10 localization constant 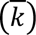 (Supplementary Fig. 9), as well as the overall size of the model domain (Supplementary Fig. 10), did not substantially alter the predicted patterns of FGF-10-induced buckling morphogenesis.

We then used our computational model to determine whether increased mesenchymal stiffness was sufficient suppress the formation of epithelial buds in response to a focal source of FGF-10 (Fig. 5A-B). Indeed, consistent with our photocrosslinking experiments, simulations with larger mesenchymal shear moduli (e.g., 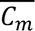 = 0.5, 1) did not produce any FGF-10-induced epithelial buds, despite similar levels of growth (*G_x_*) within the airway epithelium (Fig. 5C-D, Supplementary Movies 3-4). In these cases, epithelial growth did not produce compressive stresses sufficient to cause buckling (Fig. 5D). Instead, the magnitude of the computed stresses increased monotonically without exceeding *P_cr_* (Fig. 5D). Taken together, these computational results suggest that RB-mediated photocrosslinking can be used to modulate the mechanical properties of embryonic organs and influence epithelial patterning and morphogenesis.

**Figure 5:**
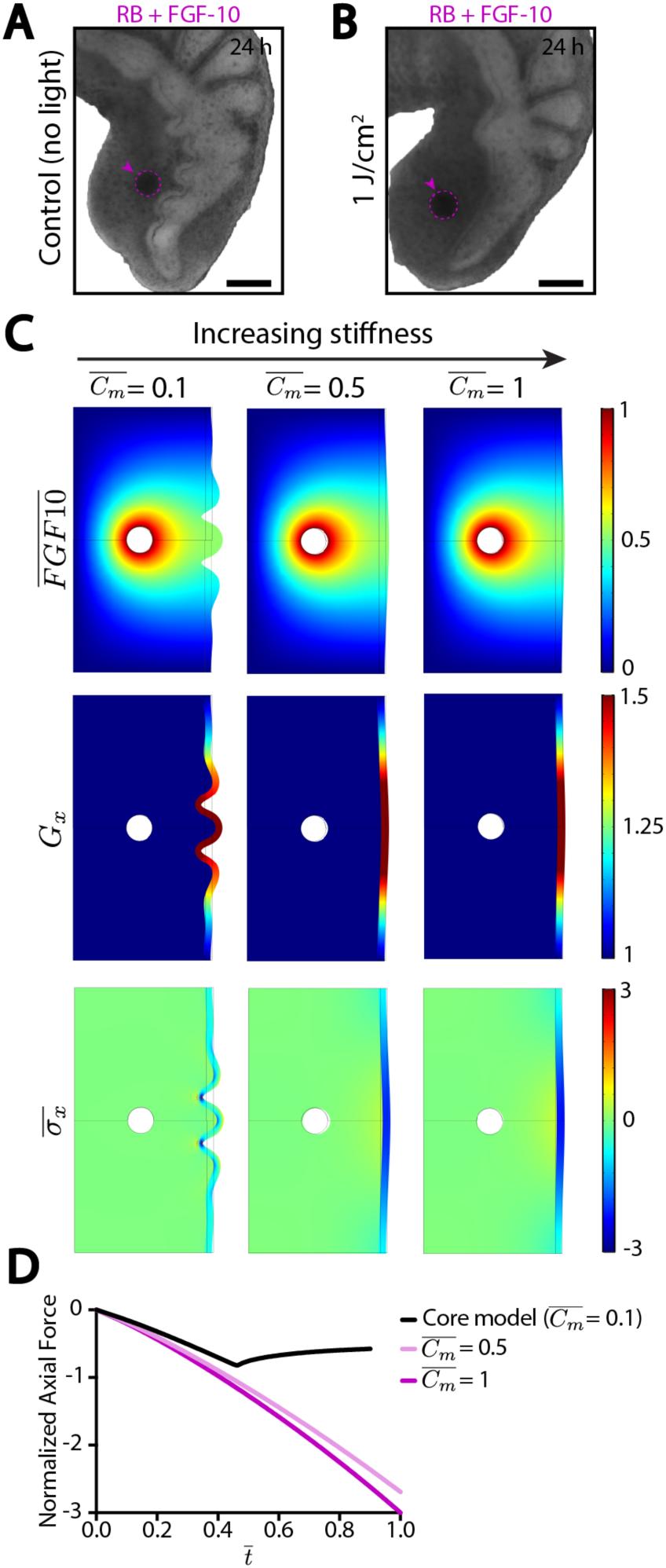
Computational model predicts that increased mesenchymal stiffness suppresses the formation of FGF-10-induced epithelial buds. (A-B) Representative bright-field images of (A) control or (B) 1 J/cm^2^ (increased stiffness) lungs showing a buckled epithelium versus suppressed buckling, respectively. Magenta dashed lines indicate location of the RB+FGF-10-loaded bead. Scale bars, 100 µm (C) Computational models with increasing mesenchymal stiffness, shown as the normalized mesenchymal shear modulus 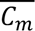. Results for normalized FGF-10 concentration 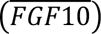, growth in the x-direction (*G_x_*), and normalized Cauchy stress 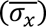 shown at the final timepoint of the models. (D) Normalized axial force versus non-dimensionalized time 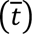 for all three models. Only the core model reaches a deflection point, interpreted as the critical load.

## Discussion

Several decades worth of experiments have investigated the mechanical properties of developing tissues ^3–6,43^. During that time, multiple studies have shown that time-dependent changes in tissue stiffness are associated with numerous developmental processes, such as gastrulation^3^, egg cleavage^5^, elongation^4^, and invagination^6^. More recently, this work has begun to investigate how differences (or gradients) in mechanical properties might influence patterns of differentiation during development^43–48^. However, these efforts have often used *in vitro* techniques with either isolated progenitor cells^49,50^ or organoid cultures^12–14^ to investigate the effect of ECM stiffness. As a result, to date, this idea has not been tested directly *in vivo*, in part owing to a lack of experimental techniques to perturb tissue stiffness within intact embryonic organs.

Here, we used RB-mediated photocrosslinking to locally modulate the mechanical properties of the pulmonary mesenchyme in cultured embryonic lungs. Photo-induced changes in tissue stiffness impacted whether or not epithelial buds formed in response to FGF-10. Importantly, treatment with RB did not negatively affect the patterns of cell proliferation and apoptosis associated with budding morphogenesis. Instead, increased mesenchymal stiffness suppressed epithelial buckling and FGF-10-induced budding events. Consistently, a computational model of the embryonic airway and surrounding pulmonary mesenchyme indicated that changes in mechanical properties within developing organs can profoundly influence epithelial morphogenesis and patterning.

Photoactivatable molecules have been previously to modulate the stiffness of optically accessible adult tissues, such as the cornea and skin^18–21^; however, our current study is the first to employ this technique to alter the mechanical properties within a developing embryonic organ. We capitalized on the capability of RB to covalently crosslink the ECM, particularly collagen fibrils^19^. In a simplified collagen gel system, significant RB-induced crosslinking was demonstrated previously using scanning electron microscopy (SEM) and produced an increase in the apparent stiffness of the collagen network ^15^, similar to the results included above (Supplementary Fig. 3). The protein-protein crosslinking caused by RB is often attributed to the production of singlet oxygen via energy transfer (type II reaction)^21^; however, molecular dynamics simulations and experimental data have also suggested that RB crosslinking in low oxygen settings might favor electron transfer (type I reaction) with positively-charged amino acids, such as arginine and lysine^16,51^. Consistent with this idea, RB crosslinking within the rabbit cornea was enhanced using arginine in an oxygen-free setting and resulted in an increased Young’s modulus as compared to controls^52^. Still, the exact mechanism by which activated RB produces protein-protein crosslinks is unclear and remains an area of active investigation. Regardless, the extent of RB-mediated crosslinking can be tuned by varying the irradiance and fluence of the activating light^19^, and these values must be carefully selected depending on the application and desired result. For instance, the ability of photosensitizers like RB to degrade ECM and cause cell death is commonly used during photodynamic therapy (PDT) to improve cancer treatment^53,54^. Here, to modify the mechanical properties of developing tissues, RB is particularly advantageous owing to its activation with green, as opposed to ultraviolet (UV) light, which is needed for riboflavin-mediated crosslinking but is more phototoxic^19^. Even so, photocrosslinking requires the tissue to be optically accessible, and this could limit its *in vivo* use within certain animal models, such as developing mouse embryos. In such instances, *ex vivo* organ culture could be combined with RB-mediated photocrosslinking, as well as genetic modulation of various ECM crosslinking enzymes, such as lysyl oxidase (LOX), or LOX-like-mediated remodeling^55^ and non-enzymatic collagen glycation^56^, to modulate tissue stiffness *in vivo*.

Here, we showed that local changes in stiffness can modulate FGF-10-induced buckling morphogenesis, and these data indicate that regional differences in mechanical properties can influence epithelial folding events, which involve a physical instability, such as gut villification, intestinal looping, cortical folding of the brain, crypt formation within the small intestine, and tooth invagination^12,13,38,57–61^. Although previous experiments using isolated airway epithelial explants in 3D gels of reconstituted basement membrane protein showed that branch formation and spacing was unaffected by differences in gel stiffness^29^, here we found that increased mesenchymal stiffness can suppress FGF-10-induced budding within intact embryonic lungs. During normal lung development, new lateral branches emerge along the length of the primary bronchus in a proximodistal fashion^25^; however, it is unclear if spatiotemporal patterns of tissue stiffness influence these branching dynamics. Still, ECM patterning has been implicated in the morphogenesis of new airways in the embryonic lung^41^, and similar mechanisms are thought to contribute to kidney^62^, mammary gland^10^, and salivary gland development^9^. FGF-10 expression is also an important regulator airway branching morphogenesis^36^, but our data suggest that focal sources of FGF-10 are not the sole determinant of the overall branching pattern. Although computational modeling has been used to predict patterns of FGF-10 expression^63^, these simulations have not yet incorporated explicitly the role of tissue mechanics. Our photocrosslinking experiments emphasize the importance of biophysical factors during airway branching morphogenesis^27,29,30,32,33,41^, as well as the interplay between growth factor signaling and mechanical forces^33,35,64^.

We also believe that RB-mediated photocrosslinking can be incorporated into dynamic studies of organ morphogenesis. For instance, the mechanism of formation for tissues that appear to undergo a physical instability, such as buckling, folding, bending, or looping^12,13,38,57–61^ can be further validated using RB photocrosslinking. Further, we propose this technique, which only impacts the mechanical properties of the ECM, can be used to parse the hypothesized mechanical inputs that either smooth muscle cells or the ECM provide to sculpt the morphogenesis of a variety of branched epithelial networks^9,10,41,62,65,66^. Although here we focused on epithelial buckling, a similar approach can be used to investigate the role of tissue mechanics in cell differentiation. Spatial differences in both gene expression encoding for an ECM-modifying enzyme as well as cells involved in maintenance of the microenvironment have been observed within the developing kidney^62^, suggesting differences in matrix stiffness may direct organogenesis. A combination of novel techniques to modulate tissue stiffness, such as RB photocrosslinking, and methods to measure *in vivo* mechanical properties will lead to an overall greater understanding of how biophysical cues direct morphogenesis and development.

## Methods

### Ex Vivo Culture of Embryonic Lungs

Fertilized White Leghorn chicken eggs were incubated in a humidified forced-draft incubator at 37°C until HH stage 26^67^. Fine forceps were used to dissect embryonic lungs under a stereomicroscope (SZX7, Olympus; Waltham, MA) in PBS supplemented with antibiotics (50 U/ml penicillin/streptomycin, Invitrogen; Carlsbad, CA). Embryonic lung explants were then cultured at the fluid-air interface for 24 hr at 37°C on Nucleopore membranes (25 mm-diameter, pore size: 8 μm) in DMEM/F12 medium (without HEPES) that had been supplemented with 5% fetal bovine serum (FBS, heat inactivated) and antibiotics (50 U/ml penicillin/streptomycin)^68^. To elicit FGF-10-induced supernumerary buds, agarose beads (100-200 mesh Affi-Gel Blue Media, Bio Rad; Hercules, CA) containing exogeneous FGF-10 (345-FG, R&D Systems; Minneapolis, MN) were implanted within the pulmonary mesenchyme along regions of the primary bronchus that do not normally branch, as previously described^37^. In some experiments, the culture medium was supplemented with 1 µM staurosporine (9953S, Cell Signaling; Danvers, MA) to induce cell apoptosis^27^. Bright-field images of cultured lung explants were captured at 0, 3, and 24 hr using a Zeiss AxioVert microscope equipped with a 5×, NA 0.15, LD A-Plan objective (Zeiss; Oberkochen, Germany).

### Photocrosslinking using Rose Bengal

RB was delivered to the embryonic lungs by introducing RB-soaked agarose beads, similar to the technique used to introduce growth-factor loaded beads as previously described^37^. Briefly, small agarose beads approximately 75-150 μm in diameter (100-200 mesh Affi-Gel Blue Media, Bio Rad; Hercules, CA) were rinsed overnight in PBS at 4°C and then allowed to soak in droplets of either 100 μg/mL recombinant human FGF-10 (345-FG, R&D Systems; Minneapolis, MN) in PBS or 100 μg/mL RB (330000, Sigma; St. Louis, MO) in DI water + 100 μg/mL rhFGF-10 in PBS for 2-3 hr at room temperature. Using a pair of fine forceps, small incisions were made within the pulmonary mesenchyme near the ventral epithelium and a single bead was placed within. The explants were then cultured for 3 hr at 37°C, as described above, to allow the RB to diffuse from the bead into the surrounding tissue. Afterwards, a subset of lungs were then exposed to 555 nm light (BioLambda; São Paulo, Brazil) to deliver an irradiance of 25 mW/cm^2^ and fluence of either 0.1 J/cm^2^, 0.5 J/cm^2^, 1 J/cm^2^, 2 J/cm^2^, 5 J/cm^2^, or 10 J/cm^2^ (control lungs were not exposed, as indicated by “no light”). After irradiation, lungs were then allowed to culture for the remainder of the 24 hr.

### Immunofluorescent staining and imaging

Cultured explants were fixed in 4% paraformaldehyde in PBS for 15 minutes, permeabilized and blocked using a solution of 10% goat serum and 0.3% Triton X-100 in PBS (PBST) for 1 hr at room temperature. To detect proliferating cells in S-phase, the Click-iT EdU Imaging Kit (Invitrogen; Carlsbad, CA) was used (explants were pulsed with EdU for 20 minutes preceding fixation). Samples were then incubated with primary antibody overnight on a shaker in 4°C. The following primary antibodies were used: anti-LCAM (1:100 dilution, 7D6, Developmental Studies Hybridoma Bank; Iowa City, IA) and anti-cleaved caspase-3 (Asp175) (1:200 dilution, 9661, Cell Signaling; Danvers, MA). Samples were washed six times for 1 hr using PBS, and then incubated with Alexa Fluor-conjugated secondary antibody overnight on a shaker in 4°C (1:200 dilution, Invitrogen; Carlsbad, CA). Stained explants were dehydrated in a graded methanol series and optically cleared using Murray’s clear. Whole-mount samples were then imaged using a Zeiss LSM 800 laser scanning confocal microscope with a 20×, NA 0.8, Plan-Apochromat objective (Zeiss; Oberkochen, Germany).

### Micromechanical testing of embryonic lung explants

Cultured explants underwent unconfined-compression testing using an ∼70 µm cylindrical fabricated microindenter with a Microsquisher (MT G2, CellScale; Waterloo, ON, Canada) to determine differences in tissue stiffness, similar to previously described^69^. The pulmonary mesenchyme of cultured explants was compressed using the cylindrical microindenter in a region of the mesenchyme near the implanted bead on each lobe. The compression test settings within the SquisherJoy software (Version 5.23, CellScale; Waterloo, ON, Canada) specified a total displacement of 500 µm over a 30s ramp. The indenter tip was positioned ∼380 µm above the top of the sample, resulting in samples undergoing ∼120 µm of displacement (exact tissue displacement was calculated in post-processing, as described below). Samples were tested a minimum of two times, and the axial force and total displacement were tracked by the front-camera and SquisherJoy software (Supplementary Fig. 2). The resulting force vs time curves were plotted and used to determine the time of contact (*t_c_*) (Supplementary Fig. 2)^69^. These curves were used to determine the tissue deflection (*δ_T_*) and the resulting force vs tissue deflection curves were plotted for each test (Supplementary Fig. 2)^69^. The two replicate curves were averaged for each sample, and sample curves were averaged across their respective treatment group to create the overall force vs tissue deflection curves (Fig. 1). Stiffness values were calculated by fitting a nonlinear curve^7^ to each individual lung’s force vs tissue deflection curve using Matlab (Version R2018A; Natick, MA) to obtain four free parameters. The tissue stiffness was then calculated as a function of tissue deflection and these parameters^7^ and averaged across treatment group to obtain a mean stiffness value and standard deviation. The stiffness value obtained at a tissue deflection of 60 µm was selected for comparison between groups (Fig. 1).

### Collagen Gel Preparation and Mechanical Testing

Collagen solutions were obtained by mixing acid-solubilized rat tail collagen I (354249, Corning; Corning, NY) with an equal volume of a neutralizing buffer (100mM HEPES solution in 2x PBS, at pH 7.3)^70^. The final concentration of collagen was adjusted to 4 mg/mL by adding adequate volumes of neutralized collagen solution and ice cold 1x PBS. This mixture was aliquoted into cylindrical PDMS molds with a diameter of 5mm and a height of 2mm, and then allowed to polymerize for 1 hour at 37°C. To achieve cross-linking, the plugs of polymerized collagen were extracted from their molds and immersed in a 100 µg/mL solution of RB for 24 hr, and then exposed to a 555 nm laser source with an irradiance of 25 mW/cm^2^ and a fluence of 0.5 J/cm^2^ (BioLambda; São Paulo, Brazil). The macroscopic properties of cylindrical collagen plugs were estimated using a microscale compression system (Cell Scale; Waterloo, ON, Canada). Using established protocols^71^, collagen gels were subjected to uniaxial unconfined compression testing by means of a 6mm x 6mm platen fixed to a microbeam of known length (57 mm), diameter (0.4064 mm) and elastic modulus (411 GPa). Five stress relaxation steps were imposed by applying a deformation equal to 2% of the unloaded gel height, followed by a 3-minute hold phase. In virtue of the nearly linear responses observed at equilibrium, the bulk mechanical behavior measured under unconfined compression was modeled using a compressible neo-Hookean strain energy density function:

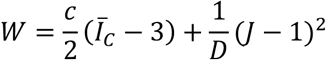

The material stiffness of the photocrosslinked collagen networks was reported as the neo-Hookean shear modulus *c* ^72^.

### Computational Modeling

A nonlinear computational model of the embryonic airway was created in COMSOL Multiphysics (Version 5.5; Burlington, MA). The epithelium was modeled as a growing layer supported by a foundation of mesenchyme. The modeled lung was assumed to behave as a hyperelastic material with a Blatz-Ko strain-energy density function, *W*:

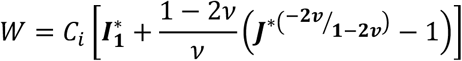

where *C*_*_ is the shear modulus of either the epithelium or the mesenchyme, 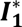 is the first elastic strain invariant, *v* is Poisson’s ratio, and ***J***^∗^ is the elastic volume ratio. The shear modulus of the epithelium (*C_e_*) was set to equal 1 while the mesenchyme (*C_m_*) was set to 0.1 (except in the increased stiffness perturbation, as noted below). Poisson’s ratio (*v*) was set to 0.49 for the entire model, making it nearly incompressible.

Growth was included by decomposing the overall deformation gradient tensor ***F*** into a component due to growth (***G***) and a component due to elastic deformation (***F***^∗^) via ***F*** = ***F***^∗^ · ***G*** ^73^. Longitudinal growth was included in the epithelium where the growth in the x-direction, ***G***_***x***_, was described by:

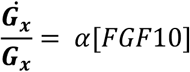

where *α* is a defined growth constant and [*FGF*10] is the concentration of modeled growth factor. The modeled mesenchyme was given a no growth condition. The Cauchy stress tensor, ***σ***, depends only on the elastic deformations and was given by:

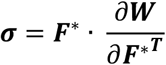

We report the non-dimensionalized Cauchy stress in the x-direction, 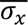, given by 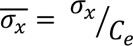. The edge of the modeled bead was given the initial condition of [*FGF*10] = 1 while the outer edge of the mesenchyme was given the boundary condition of [*FGF*10] = 0, to recapitulate the media sink (Supplementary Fig. 5). The outer edge (luminal/apical surface) of the epithelium had a zero flux condition, mimicking the barrier function it performs (Supplementary Fig. 5). Diffusion of the modeled growth factor was assumed to follow Fick’s Second Law of Diffusion. We assumed one-dimensional, steady-state diffusion, which simplifies to the following:

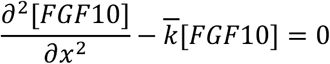

where 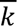 is the non-dimensionalized localization parameter given by 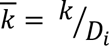. Both diffusion constants for the epithelium (*D_e_*) and mesenchyme (*D_m_*) were set to equal 1. The non-dimensionalized localization parameter 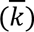 was set to equal 1 in the core model (unless otherwise noted in the varying localization parameter models). Additionally, to reduce the computational load and computing time, the geometry was halved symmetrically along the y-axis midline and results were mirrored. A small positive point force in the y-direction was added in the middle of the epithelium to disrupt equilibrium. Except for the outer edge of the epithelial layer in the x-direction, all other outer edges were constrained using rollers (Supplementary Fig. 5).

The model underwent various perturbations to test its robustness as well as the biophysical mechanisms from experimental data, including varying 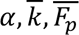 as well as the location of the growth factor source, mesh size, and domain size (Supplementary Fig. 6-10). The photoactivatable crosslinking and tissue stiffening induced by RB treatment was recapitulated in the model by increasing the shear modulus of the mesenchyme while keeping all other aspects the same.

### Statistics

Data represent means ± standard deviation for at least three independent experiments. Statistical comparisons were made using either a Student’s t-test, or a one- or two-way ANOVA, followed by a Tukey post-hoc test, with p-values as specified in the figure legends.

## Acknowledgments

This work was supported by grants from the National Institutes of Health (R01HL145147 and R01HL133163).

## Author Contributions

Conceptualization: K.E.P., V.D.V.; Formal analysis: K.E.P., P.R., A.K.; Investigation: K.E.P., P.R., A.K; Writing - original draft: K.E.P.; Writing - review & editing: K.E.P., A.K., J.P.G., G.O., J.F., V.D.V.; Supervision: G.O., J.F., V.D.V.; Project administration: V.D.V.; Funding acquisition: V.D.V.

