## Supplementary Material for "Photo-induced changes in tissue stiffness alter epithelial budding morphogenesis in the embryonic lung"

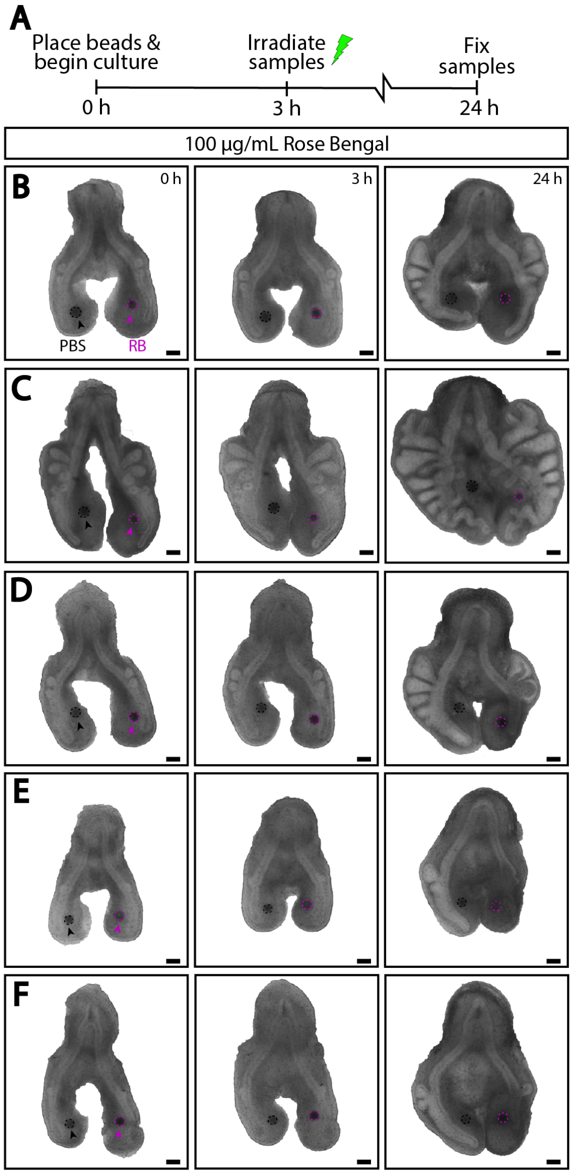


**Supplementary Fig. 1: Troubleshooting Rose Bengal treatment and photocrosslinking.** (A) Experimental timeline. Samples were allowed to culture for 3 h, then were briefly exposed to green light to deliver an irradiance of 25 mW/cm$^2$ and varying fluence values of 0.5 J/cm$^2$, 1 J/cm$^2$, 2 J/cm$^2$, 5 J/cm$^2$, or 10 J/cm$^2$. Lungs were allowed to culture the remainder of the 24 h and then fixed. (B-F) Representative bright-field images of lungs cultured with either PBS- (left lobe, black arrow and dashed lines) or RB-loaded beads (right lobe, magenta arrow and dashed lines). Darkened, unhealthy tissue is seen within lungs exposed to the higher fluence values (≥2 J/cm$^2$) (E-F). Scale bars, 200 $\mu$m.

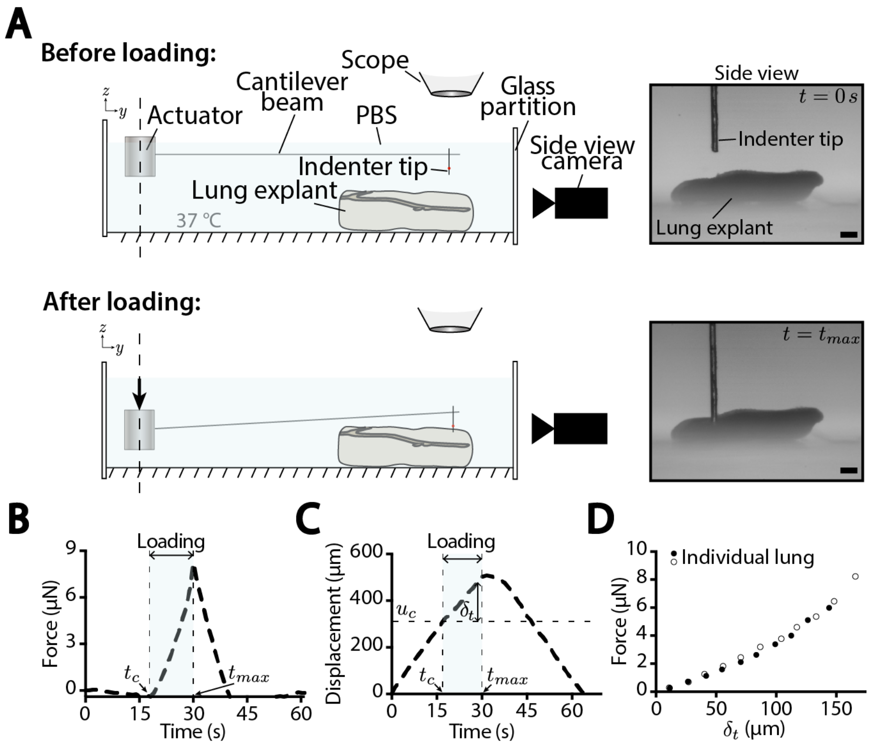


**Supplementary Fig. 2: Microindentation mechanical testing and analysis.** (A) Schematic of microindentation mechanical testing of lung explants using the Microsquisher (CellScale). Representative side view bright-field images shown before loading at $t = 0\ s$ and at the end of loading at $t = t_{max}$. Scale bars, 100 $\mu$m. (B) Representative force versus time curve for a tested explant used to determine the time of contact $(t_c)$. Shaded region shows the loading regime, taken as the time between the time of contact $(t_c)$ and time of maximum force $(t_{max})$. (C) Representative total displacement versus time curve for a tested explant used to determine tissue deflection $(\delta_t)$, calculated as the difference between the displacement during loading $(u)$ and displacement at the time of contact $(u_c)$. (D) Representative force versus tissue deflection $(\delta_t)$ for an individual lung explant mechanically tested two times.

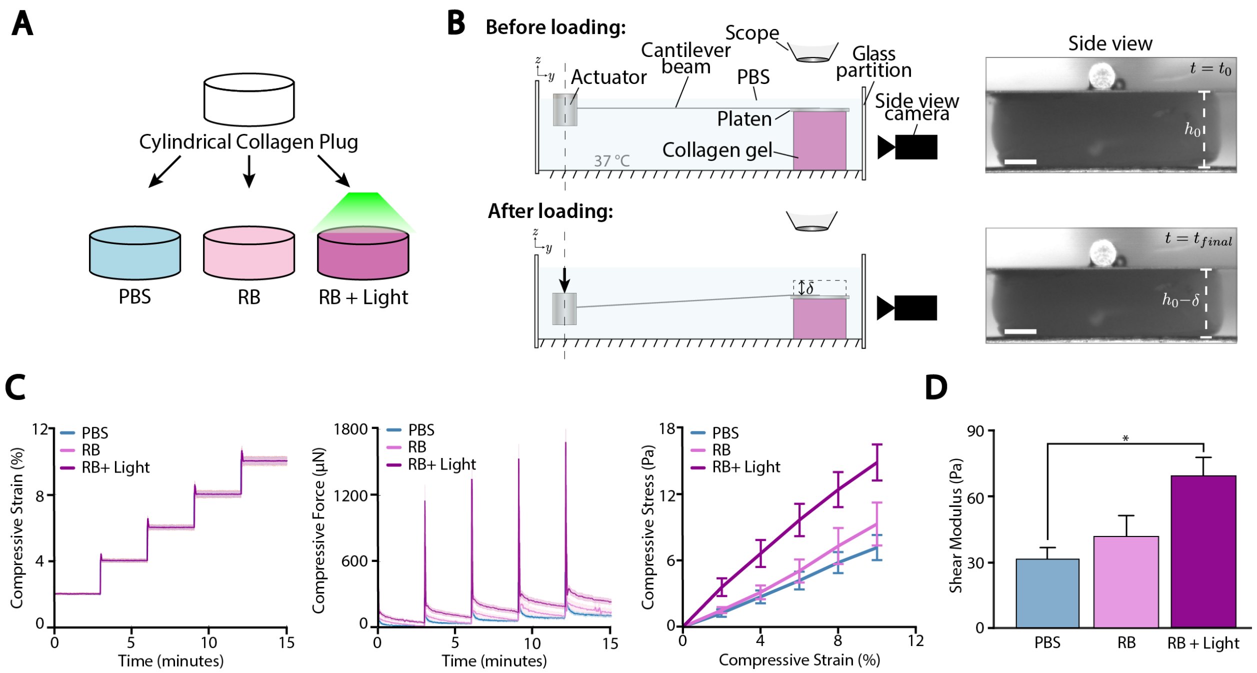


**Supplementary Fig. 3: Photocrosslinking and stiffening in collagen gels.** (A) Cylindrical collagen plugs were divided into three groups, based on exposure to one of three conditions: immersion in 1x PBS for 24 hours (PBS), immersion in a 100 $\mu$g/mL solution of Rose Bengal for 24 hours without exposure to light (RB), or immersion in a 100 $\mu$g/mL solution of Rose Bengal for 24 hours, followed by exposure to a laser source with irradiance of 25 mW/cm$^2$ and fluence of 0.5 J/cm$^2$ (RB+Light). (B) Schematic illustration of the unconfined compression experiment used to characterize collagen mechanics. Displacement of a platen ($\delta$) connected to a linear actuator via a cantilever beam of known stiffness was used to quantify axial forces generated by the collagen plugs undergoing compression. At the same time, a camera was used to image the reference ($h_0$) and deformed ($h_0 - \delta$) heights of the collagen plugs and quantify the compressive strain as $\varepsilon = 1 - (h_0 - \delta)/h_0$. Scale bars, 500 $\mu$m. (C-D) Unconfined compression was performed in 5 incremental steps, each with a magnitude equal to 2% of the original height of the sample, interspersed with 3-minute long hold phases. Averaged plots show the compressive force versus time for the different treatment groups. For each sample, the equilibrium force-displacement data is converted into a compressive stress versus compressive strain plot and fitted to a neo-Hookean material model[72] to quantify the shear modulus (D). The modulus increases significantly upon photo-crosslinking of collagen with Rose Bengal. Data are presented as mean ± SEM. Statistical significance was determined using a one-way ANOVA with a Tukey post-hoc test. (PBS: n=5; RB: n=5; RB+Light: n=9; $^*p < 0.05$).

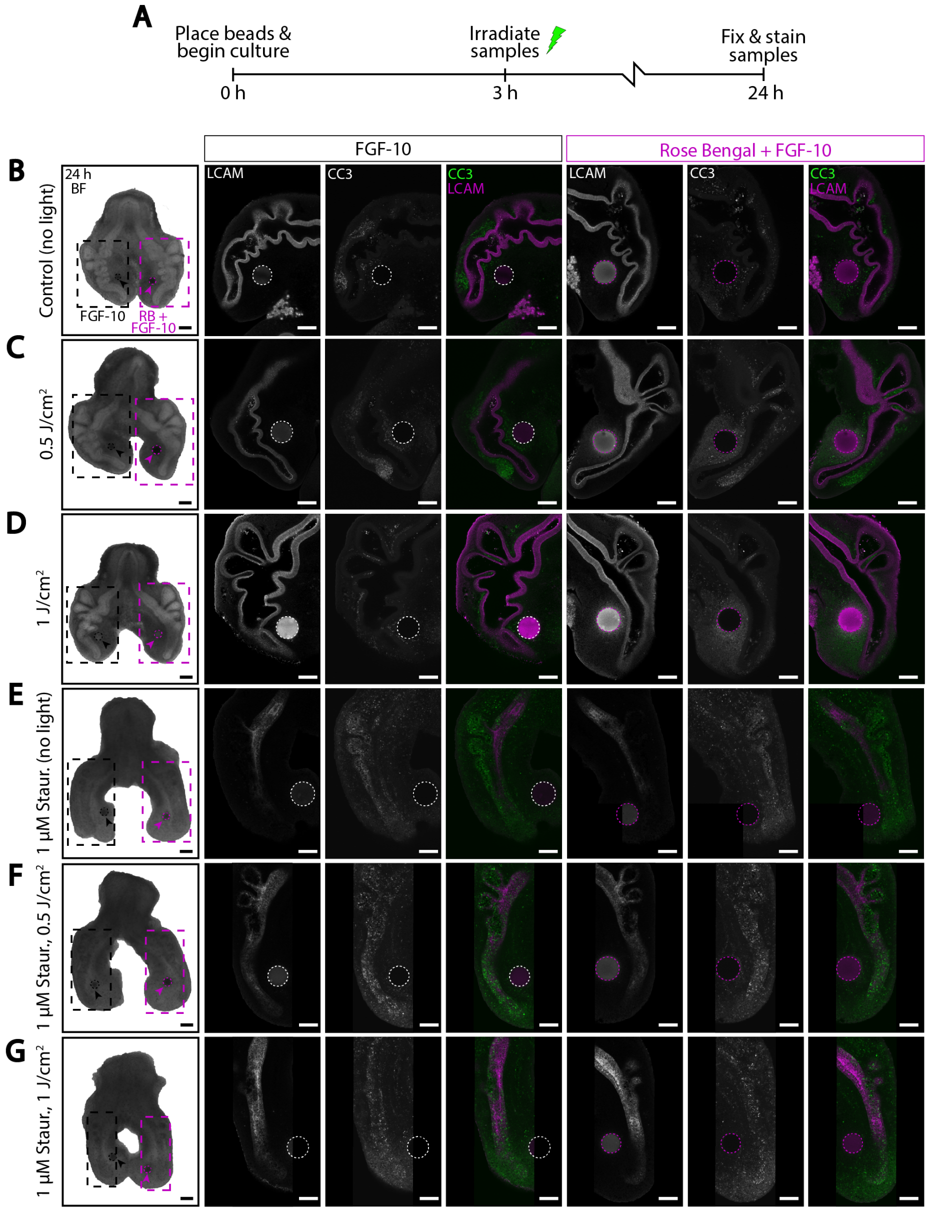


**Supplementary Fig. 4: CC3 signal elevated within staurosporine-treated lungs compared to untreated.** (A) Experimental timeline. Samples were allowed to culture for 3 h, then some were briefly exposed to green light to deliver an irradiance of 25 mW/cm$^2$ and varying fluence values of either 0.5 J/cm$^2$ or 1 J/cm$^2$. Lungs were allowed to culture the remainder of the 24 h after irradiation and then fixed and stained. Some explants were cultured with 1 $\mu$M staurosporine (staur.), an activator of cell apoptosis. (B-G) Representative bright-field images of lung explants. Insets show representative confocal images of lungs stained for CC3 (cell apoptosis marker) and LCAM (E-cadherin marker) immunofluorescence along with the merged image. (E-G) Treatment with 1 $\mu$M staurosporine resulted in dark, unviable lungs that did not form new epithelial branches and showed increased CC3 signal within the epithelium as compared to lungs not treated with staurosporine (A-D). Black or white dashed lines show the location of FGF-10-soaked beads. Magenta dashed lines show the location of RB+FGF-10-soaked beads. Bright-field images scale bars, 200 $\mu$m. Confocal images scale bars, 100 $\mu$m.

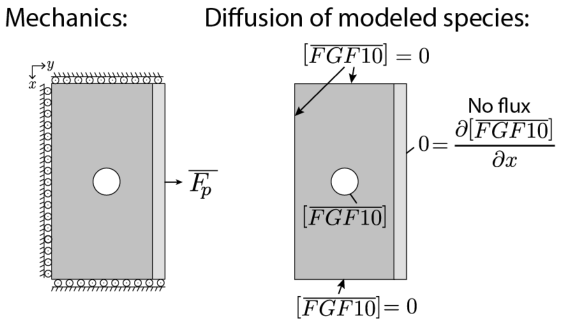


**Supplementary Fig. 5: Computational model boundary conditions.** Rollers were used to constrain the motion of the outer edges of the mesenchyme to one direction. A small non-dimensionalized point force $\overline{F_p}$ (~$10^{-4}$ $\mu$N) was used to perturb the epithelium from an unstable equilibrium. The normalized concentration of FGF-10 ($[\overline{FGF10}]$) was specified at the bead boundary and set to 0 at the outer edges of the mesenchyme. A no flux boundary condition was specified at the outer edge of the epithelium.

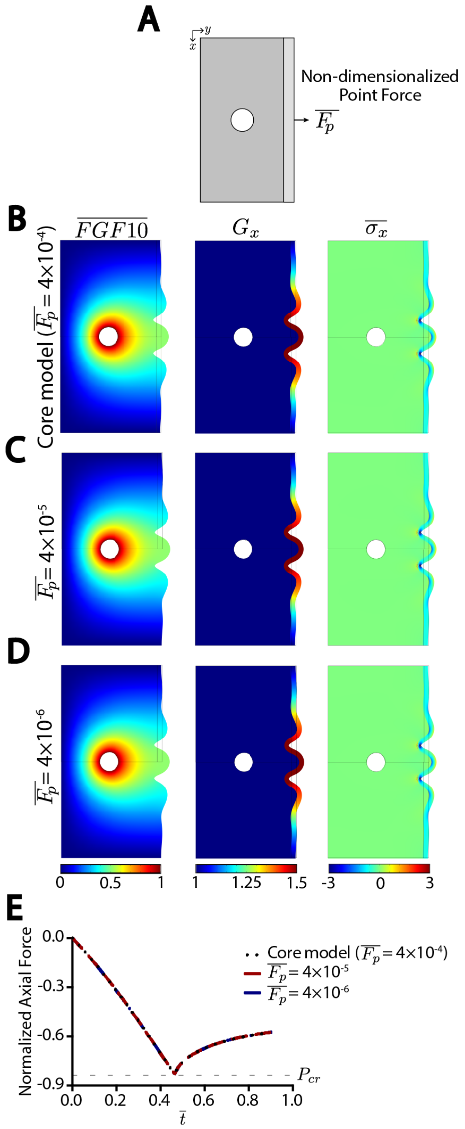


**Supplementary Fig. 6: Effect of varying magnitude of non-dimensionalized point force in computational model.** (A) A small non-dimensionalized point force $\overline{F_p}$ was used to perturb the epithelium from an unstable equilibrium and its magnitude was varied to test the robustness of the model. (B-D) Results for normalized FGF-10 concentration $(\overline{FGF10})$, growth in the x-direction $(G_x)$, and normalized Cauchy stress $(\overline{\sigma_x})$ shown at the final timepoint of the models. (E) Normalized axial force versus non-dimensionalized time $(\bar{t})$ for all three models. All curves overlap and reach a deflection point, interpreted as the critical load $(P_{cr})$ and denoted by the horizontal dashed line.

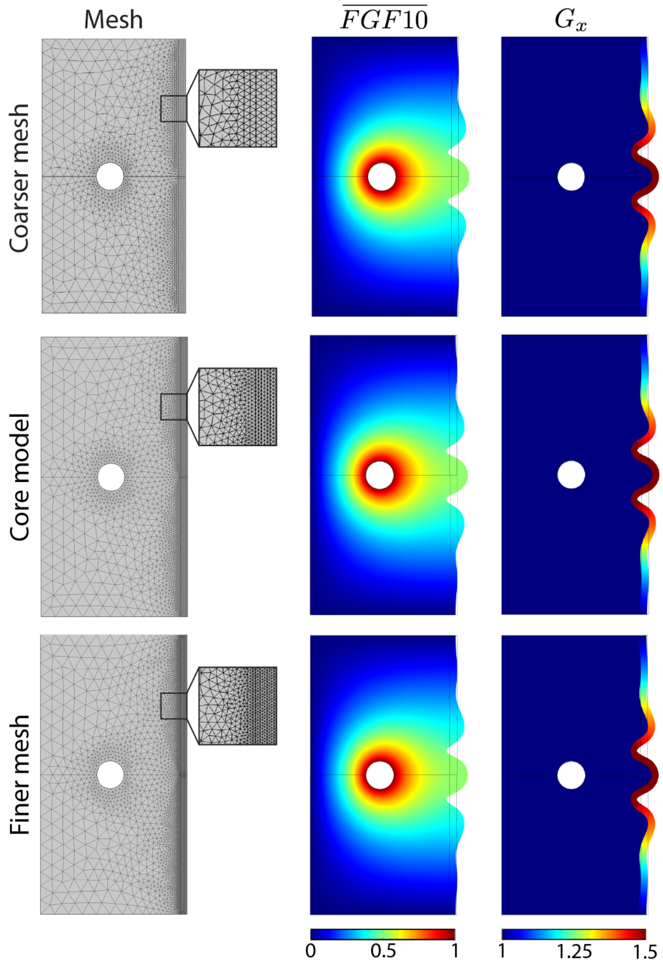


**Supplementary Fig. 7: Effect of varying size of mesh in computational model.** The maximum element size within the epithelium was increased or decreased to create a coarser or finer mesh, respectively. Results for normalized FGF-10 concentration $(\overline{FGF10})$ and growth in the x-direction $(G_x)$ shown at the final timepoint of the models.

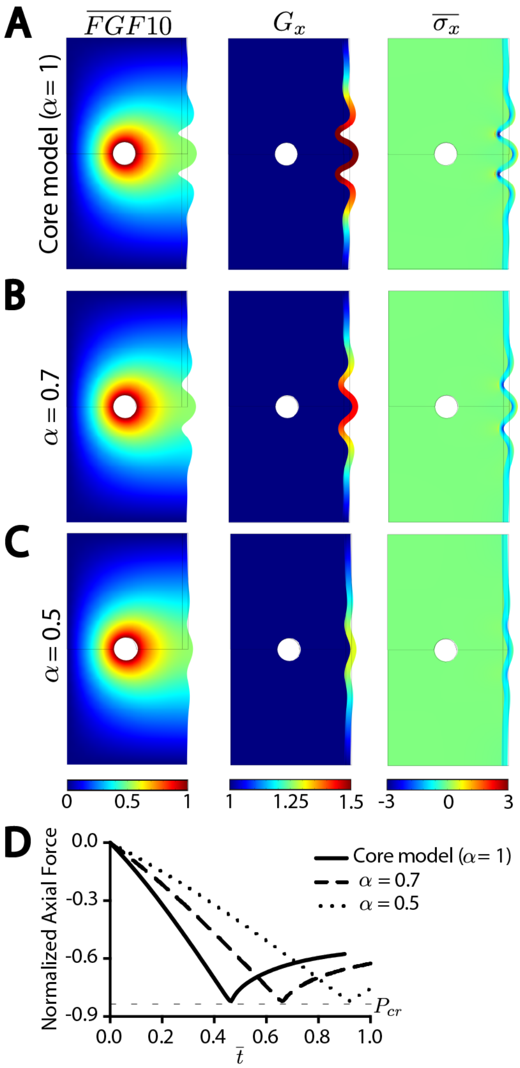


**Supplementary Fig. 8: Effect of varying growth constant in computational model.** (A-C) The growth constant $(\alpha)$ was decreased. Results for normalized FGF-10 concentration $(\overline{FGF10})$, growth in the x-direction $(G_x)$, and normalized Cauchy stress $(\overline{\sigma_x})$ shown at the final timepoint of the models. (E) Normalized axial force versus non-dimensionalized time $(\bar{t})$ for all three models. As $\alpha$ increases, the time at with the curve reaches a deflection point, interpreted as the critical load $(P_{cr})$ and denoted by the horizontal dashed line, shifts to the right.

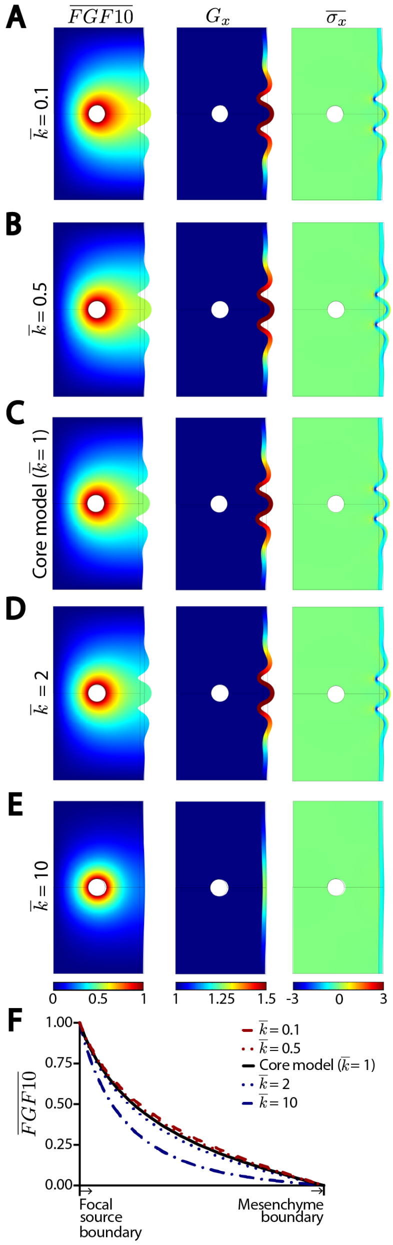


**Supplementary Fig. 9: Effect of varying localization constant in computational model.** (A-E) The normalized localization constant $(\overline{k})$ was varied to tune the focalness of FGF-10 diffusion. Results for normalized FGF-10 concentration $(\overline{FGF10})$, growth in the x-direction $(G_x)$, and normalized Cauchy stress $(\overline{\sigma_x})$ shown at the final timepoint of the models. (F) Changes in $\overline{FGF10}$ is plotted starting at the boundary edge of the focal source along the x-direction and ending at the mesenchyme boundary for all models.

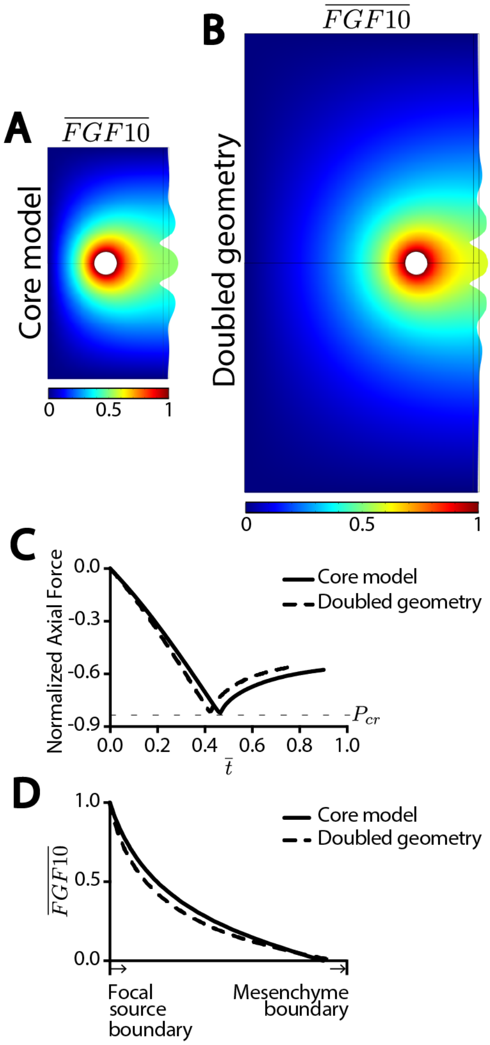


**Supplementary Fig. 10: Effect of varying domain size in computational model.** (A-B) The dimensions of the mesenchyme in the core model (A) are doubled (B). Results for normalized FGF-10 concentration $(\overline{FGF10})$ and the deformation are shown for the final timepoint. (C) Normalized axial force versus non-dimensionalized time $(\bar{t})$ for both models. The curves are similar and reach a deflection point, interpreted as the critical load $(P_{cr})$ and denoted by the horizontal dashed line. (D) Changes in $\overline{FGF10}$ is plotted starting at the edge of the focal source along the x-direction and ending at the mesenchyme boundary for all models. The curve for the doubled geometry model was normalized to the core model geometry to allow for comparisons.

**Supplementary Movie 1:** Modeled diffusion of FGF-10 concentration within the core model and resulting deformations.

**Supplementary Movie 2:** Modeled axial growth within the core model and resulting deformations.

**Supplementary Movie 3:** Modeled axial growth within the increased stiffness model ($\overline{C_m} = 0.5$) and resulting deformations.

**Supplementary Movie 4:** Modeled axial growth within the increased stiffness model ($\overline{C_m} = 1$) and resulting deformations.
